# Can a Simple Phenomenological Model Explain the Mechanics of Eccentric Contractions?

**DOI:** 10.1101/2023.03.12.532315

**Authors:** Sang Hoon Yeo, Walter Herzog

## Abstract

Notwithstanding decades of research, predictions from the existing muscle models on eccentric contraction remain elusive. This study aimed to test the possibility that a relatively simple phenomenological muscle model, based on an adapted Hill-type muscle model with some novel assumptions, can capture the mechanics of eccentric contractions. The result of the simulations, although preliminary, support the potential of the proposed model in predicting the mechanics of eccentric contraction.

## Introduction

Unlike its odd name, eccentric contraction—the lengthening of an active muscle due to excessive external force—is one of the common contractile scenarios that muscles undergo during normal everyday functioning. The classic Hill-type Muscle Model (HMM) [1], a phenomenological representation of the cross-bridge model, is widely used in upper-layer studies, such as musculoskeletal modelling and simulation, but the performance of the HMM in predicting eccentric contractions has not been satisfactory. Alternative models have been proposed to expand the explanatory scope of the cross-bridge model to include eccentric contractions [2, 3], but these models suffer from high parametric complexity, which makes them unsuitable for practical applications in upper-layer studies.

For this reason, our study was focused on testing if a simple phenomenological model, whose complexity is similar to that of the HMM, can predict representative mechanics of eccentric contractions. In contrast to most previous models, we assumed that (1) the Force-Velocity (FV) relationship is multiplied with the total (i.e. active plus passive) Force-Length (FL) function, while FV in previous models had been assumed to be multiplied with the active FL relationship only, and that (2) a certain length of the Passive-elastic Element (PE) is clamped upon activation and the stiffness of the passive FL curve is enhanced around the clamped length.

## Methods

We used isokinetic force profiles of in-situ cat soleus. Details of the experimental methods used for data collection can be found in Lee et al [4]. Data analysis was performed in Matlab.

## Result and Discussion

We focused on the force enhancement patterns during and after eccentric contractions: dynamic Force Enhancement (dFE) and residual Force Enhancement (rFE) highlighted in Fig-1 right (dotted boxes). We first focused on the temporal profiles of dFE that show an interesting transition from convex to concave (Fig-1 right, inset) as length increases. When these patterns are re-plotted in the force-length space by combining length (Fig-1 left) and force (Fig-1 right) profiles, we found that dFE patterns (Fig-2 left, orange curves) become similar to the total FL curve (Fig-2 left, blue curve).

**Figure 1.**
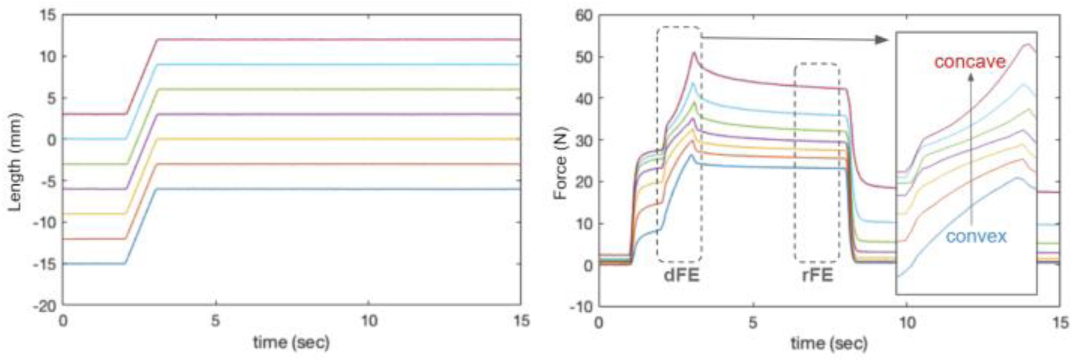
Residual and dynamic force enhancement of cat soleus [4]

**Figure 2.**
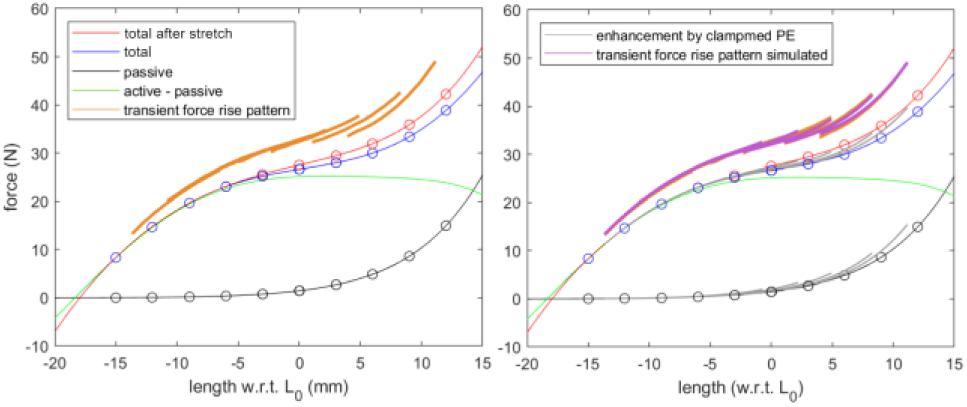
dFE patterns (data and prediction) plotted with FL curves

Inspired by this similarity, we assumed that dFE patterns originate from the total FL relationship multiplied with the viscosity determined by the FV relationship. For constant speed isokinetic stretches, this viscosity is assumed to be constant. Next, we assumed that rFE (Fig-2, red circles compared to blue circles) occurs due to the enhancement of the PE by clamping (Fig-2 right, grey branches from the black curve). Linking these assumptions, we finally hypothesized that dFE patterns are produced by the multiplication of the total FL curve with the enhanced PE (Fig-2 right, grey branches from the blue curve) and the viscosity determined by the FV relationship.

Simulated dFE patterns using our model are shown in Fig-2 (right, purple curves). Our suggested model shows an excellent performance in predicting the pattern of dFE (*R^2^ = 0.960*). Note that the simulation has only one free parameter, which is the constant FV value.

## Conclusions

This preliminary study shows the potential of the presented simple phenomenological model in predicting the mechanics of eccentric contraction. Further validation of the proposed model under varying mechanical scenarios are ongoing.

## Acknowledgments

This work is funded by BBSRC (BB/S003762/1).

